# BDNF Overexpression in the Prelimbic Cortex Does Not Reduce Anxiety- and Depression-like Behavior in Serotonin Knockout Rats

**DOI:** 10.1101/2020.07.01.180604

**Authors:** Danielle M. Diniz, Kari Bosch, Francesca Calabrese, Paola Brivio, Marco A. Riva, Joanes Grandjean, Judith R. Homberg

**Affiliations:** Department of Cognitive Neuroscience, Centre for Neuroscience, Donders Institute for Brain, Cognition, and Behaviour, Radboud University Nijmegen Medical Centre, Nijmegen, the Netherlands; Department of Pharmacological and Biomolecular Sciences, Universita’ degli Studi di Milano, Italy; Department of Radiology and Nuclear Medicine, Donders Institute, Radboud University Medical Centre, Nijmegen, The Netherlands

**Keywords:** BDNF, Serotonin knockout rats, prefrontal cortex, depression, anxiety

## Abstract

Depressive disorders are one of the leading causes of non-fatal health loss in the last decade. Adding to the burden, the available treatments not always properly work for some individuals. There is, therefore, a constant effort from clinical and preclinical studies to bring forward a better understanding of the disease and look for novel alternative therapies. Two target systems very well explored are the serotonin and the brain-derived neurotrophic factor (BDNF) systems. Selective serotonin reuptake inhibitors (SSRIs), a commonly used class of antidepressants, target the serotonin transporter (SERT) and increase serotonin levels, which in turn also leads to an increase in BDNF. A rat model lacking SERT (SERT knockout) has been a useful tool to study the interplay between serotonin and BDNF. SERT^−/−^ rats present increased extracellular levels of serotonin, yet BDNF levels are decreased, especially in the prefrontal cortex (PFC) and hippocampus. The animals further display anxiety- and depression-like behavior. Therefore, BDNF might mediate the phenotype expressed by the SERT^−/−^ rats. In this study, we sought to investigate whether overexpression of BDNF in the brain of SERT^−/−^ rats would rescue its anxious and depressive-like behavior. Through stereotaxic surgery, SERT^−/−^ and wild-type (WT) rats received BDNF or GFP lentivirus microinfusions into the prelimbic cortex subregion of the mPFC and were submitted to the sucrose consumption, open field test, and forced swim tests. Additionally, we measured hypothalamus-pituitary-adrenal (HPA)-axis reactivity. The results revealed that SERT^−/−^ rats presented decreased sucrose intake, decreased locomotor activity, and increased escape-oriented behavior in the forced swim test compared to WT rats. BDNF upregulation in WT rats caused alterations in the HPA-axis function, resulting in elevated basal plasma corticosterone levels and decreased plasma corticosterone upon stress. In conclusion, BDNF overexpression in the PrL, in general, did not rescue SERT^−/−^ rats from its depression- and anxiety-like behavior, and in WT animals, it caused a malfunction in the HPA-axis.

## Introduction

All around the world, individuals are experiencing the damaging symptoms caused by depressive disorders, such as depressed mood, loss of interest or pleasure in nearly all activities (anhedonia), appetite and sleep disturbances, fatigue, and loos of energy. These symptoms usually lead to significant impairment in the achievement of essential tasks at home, work, or school. Additionally, depressive patients can manifest feelings of worthlessness and inappropriate guilt resulting in suicidal thoughts or actually in suicidal attempts (Morris et al., 2017). The rate of individuals suffering from depression is very high; by 2015, it had reached 322 million worldwide (WHO 2017). Thereby, depression is among the leading causes of non-fatal health loss for nearly the last three decades (GBD 2017 Disease and Injury Incidence and Prevalence Collaborators, 2018). Moreover, although many of the treatments for depression have been shown to work effectively, these treatments do not work for all individuals, and commonly after a successful treatment, many patients relapse (Cuijpers, 2017). Furthermore, despite the socio-economic impact led by this disorder, the causes leading to depression are still not fully understood. The etiology of depression is attributed to a complex interaction between environmental factors and genetic vulnerability, but specific genes have not yet been found, making it challenging to comprehend the mechanistic causes of such complex disease (Nestler et al., 2002).

Clinical and preclinical studies are attempting to elucidate the pathophysiological processes underlying depression. These efforts brought to light, for instance, the serotonin hypothesis of depression in the early 1960s, which is based on the observation that antidepressant drugs can increase the concentration of monoamines, especially serotonin, in the brain (Coppen, 1967). However, although it has been demonstrated that increasing serotonin levels using, for example, selective serotonin reuptake inhibitors (SSRIs), leads to relieve in depressive symptoms, inconclusive and inconsistent studies have shown that the serotonin hypothesis seems to be too simplistic to explain the mechanisms by which this mood disorder develops in some individuals (Thompson et al., 2015). One particular observation involves the role of the serotonin reuptake transporter (SERT), which is responsible for regulating extracellular serotonin levels, and it is the target of SSRIs. Genetic down-regulation of SERT leads to increased central levels of serotonin, reproducing, thereby, the effects of SSRIs. However, it is well established that SERT downregulation is also associated with anxious and depressive phenotypes. Therefore, although antidepressants increase serotonin levels, other systems might be involved in the antidepressant response. The SERT knockout (SERT^−/−^) rats, for example, are characterized by a complete lack of SERT and increased extracellular serotonin levels (Homberg et al., 2007), yet show anxious and depressive-like phenotypes. This animal model presented anhedonia-like behavior in the sucrose preference test, increased immobility in the forced swim test, and decreased time spent in the central part of the open field, indicating that anxiety levels and depressive-like behavior are increased (Olivier et al., 2008). Additionally, SERT^−/−^ rats displayed increased levels of basal plasma corticosterone (CORT) levels under control conditions, showing altered basal hypothalamic-pituitary-adrenal axis (HPA-axis) activity (van der Doelen et al., 2014).

Taking into consideration that simply increased serotonin levels does not necessarily lead to amelioration of depressive behavioral phenotypes, further understanding of the molecular basis of depressive disorders started to be explored in light of another hypothesis for the origin of depression, namely the neurotrophic hypothesis of depression (Duman and Monteggia, 2006). Neurotrophins have a crucial role in the synaptic maturation, neuronal growth, and synaptic plasticity both during development and adulthood (Autry and Monteggia, 2012). Impaired production, release, and/or action of this class of signaling molecules is believed to have a direct association with depression (Duman and Monteggia, 2006). Among the neurotrophin family are the neurotrophins 3 and 4/5, nerve growth factor (NGF), and the brain-derived neurotrophic factor (BDNF) (Huang and Reichardt, 2001), of which BDNF is the most abundant and one of the most investigated neurotrophins.

Several studies support the neurotrophin hypothesis of depression and point to the involvement of BDNF in the physiopathology of this disorder. Most of the clinical studies reported a reduction in BDNF protein levels in the serum of depressive individuals. These studies showed that there is a direct correlation between antidepressant treatment and an increase in peripheral BDNF protein levels of treated patients, while untreated individuals present decreased levels of BDNF protein (Fernandes et al., 2015; Polyakova et al., 2015; Sen et al., 2008). Studies also reported abnormal mRNA BDNF or TrkB expression in the hippocampus and prefrontal cortex post-mortem tissue of suicidal patients with a previous record of major depression (Dwivedi et al., 2003).

Noteworthy is the observation that, while antidepressant treatment induces increases in BDNF levels, the genetic manipulation of the SERT in rats causes a decrease in BDNF levels. As mentioned above, although inherited SERT downregulation in SERT^−/−^ rats is associated with constitutive increased levels of serotonin, these animals present anxiety- and depression-like behavior (Homberg et al., 2014). Moreover, in agreement with the neurotrophic hypothesis of depression, it was shown that SERT^−/−^ rats present, under basal conditions, downregulation of BDNF mRNA and protein levels in the hippocampus and prefrontal cortex (Molteni et al. 2010; Calabrese et al. 2013). Further, total BDNF mRNA levels (exon IX) were significantly downregulated and the reduction of BDNF gene expression observed in the prefrontal cortex of SERT^−/−^ rats was shown to be due, at least in part, to epigenetic changes affecting the promoter regions of exons IV and VI (Molteni et al., 2010). Therefore, while pharmacological increase of serotonin through SSRIs leads to an increase in BDNF levels, genetic deletion of the SERT leads to likewise increased serotonin levels but decreased in BDNF levels.

BDNF presents a complex gene structure; its regulation occurs at transcriptional, translational, and post-translational levels. The human gene presents 11 different exons regulated by nine promoters (Pruunsild et al., 2007), and the rodent gene consists of nine distinct exons with eight 5’ untranslated exons and one protein-coding 3’ exon (Aid et al., 2007). The multiple BDNF exons generate a wide diversity of BDNF transcripts that differentially control BDNF protein expression in an activity-dependent and tissue-specific manner (Autry and Monteggia, 2012; Mercado et al., 2017; Miranda et al., 2019). The BDNF mature protein is subject to post-translational modifications. It is synthesized as its precursor preproBDNF in the endoplasmic reticulum (ER), where the pre-domain is cleaved generating proBDNF; proBDNF is then transferred to the Golgi apparatus to be sorted into secretory vesicles (Lessmann et al., 2003). Extracellular or co-released endopeptidases are responsible for removing the pro-domain, which can happen in different stages following the secretion of the pro-protein (Leßmann and Brigadski, 2009). Interestingly, not only the mature form of BDNF has a cellular function. While mature BDNF has an affinity for the tropomyosin-related kinase receptor TrkB receptor promoting synaptic plasticity, proBDNF has a preference for the p75NTR receptor, which activates the pathway for cellular apoptosis (Teng et al., 2005).

Different brain regions play a role in the physiopathology of depression, with the prefrontal cortex being one of them. It has been demonstrated that major depressive disorder, for example, is associated with structural and functional brain imaging changes, including reduced brain volume and activity in the PFC (Schulz and Arora, 2015). These structural changes in depressed patients have been confirmed in post-mortem studies demonstrating a reduction in neurons and glial loss in the PFC, which is accompanied by a reduction in BDNF in this brain area (Duman and Monteggia, 2006; Krishnan and Nestler, 2008). In fact, the prefrontal cortex (PFC) is a well-known brain region responsible for processes such as cognitive, motivational, and emotional regulation (Heinz et al., 2005; Pitts et al., 2016). Interestingly, the prelimbic cortex (PrL), a subdivision of the medial prefrontal cortex (mPFC), primarily projects to limbic regions, including the nucleus accumbens and the basolateral amygdala showing not only a clear connection with the reward pathway, but also involvement with the regulation of behavioral responses to stress (Choi et al., 2012; Patel et al., 2019; Vertes, 2004).

Given the reduced levels of BDNF in the PFC of SERT^−/−^ rats (Calabrese et al., 2013; Molteni et al., 2010) and the role of BDNF in supporting neuronal plasticity that is particularly affected in depressive disorders (Miranda et al., 2019), we sought to investigate whether BDNF gene overexpression can rescue the anxiety- and depression-like behavior of these rats. We selected the PrL as a target due to its connectivity with the reward pathway, which is affected in depression and other mood disorders (Nestler and Carlezon, 2006; Vertes, 2004). For gene overexpression, the BDNF exon IV was chosen because notably, this transcript is downregulated in the mPFC of SERT^−/−^ rats (Calabrese et al., 2013).

## Material and Methods

### Animals

SERT^−/−^ rats (Slc6a41Hubr) were generated by N-ethyl-N-nitrosourea (ENU)-induced mutagenesis on a Wistar background (Smits et al., 2006). SERT^−/−^ rats were derived from crossing heterozygous 5-HT transporter knockout (SERT^+/−^) rats that were outcrossed for at least 15 generations with wild-type Wistar^CrI:WI^ rats obtained from Charles River Laboratories (Horst, the Netherlands). Ear punches were taken at the age of 21 days for genotyping, which was done by LGC (Hoddesdon, United Kingdom). SERT^+/+^ rats were used to check BDNF virus overexpression in naïve animals. For the behavioral experiments, due to breeding difficulties, we didn’t achieve the required number of SERT^+/+^ rats from the nests. Therefore, we used SERT^−/−^ rats and wild-type Wistar^CrI:WI^ rats (WT rats) from Charles River (Horst, the Netherlands) as behavioral wild-type controls (see experimental design in figure 1). All animals were housed in temperature-controlled rooms (21 °C) with standard 12/12-h day/night-cycle (lights on at 7:00 am) and food and water available ad libitum. 5-7 days before surgery, animals were socially housed in individually ventilated (IVC) cages for habituation. After surgery, animals were separately housed in the IVC cages until the end of the sucrose preference test, thereafter the animals we socially housed again and kept under the same temperature and day/night-cycle throughout the entire experiment. All experiments were approved by the Committee for Animal Experiments of the Radboud University Nijmegen Medical Centre, Nijmegen, the Netherlands, and all efforts were made to minimize animal suffering and to reduce the number of animals used.

**Figure 1.**
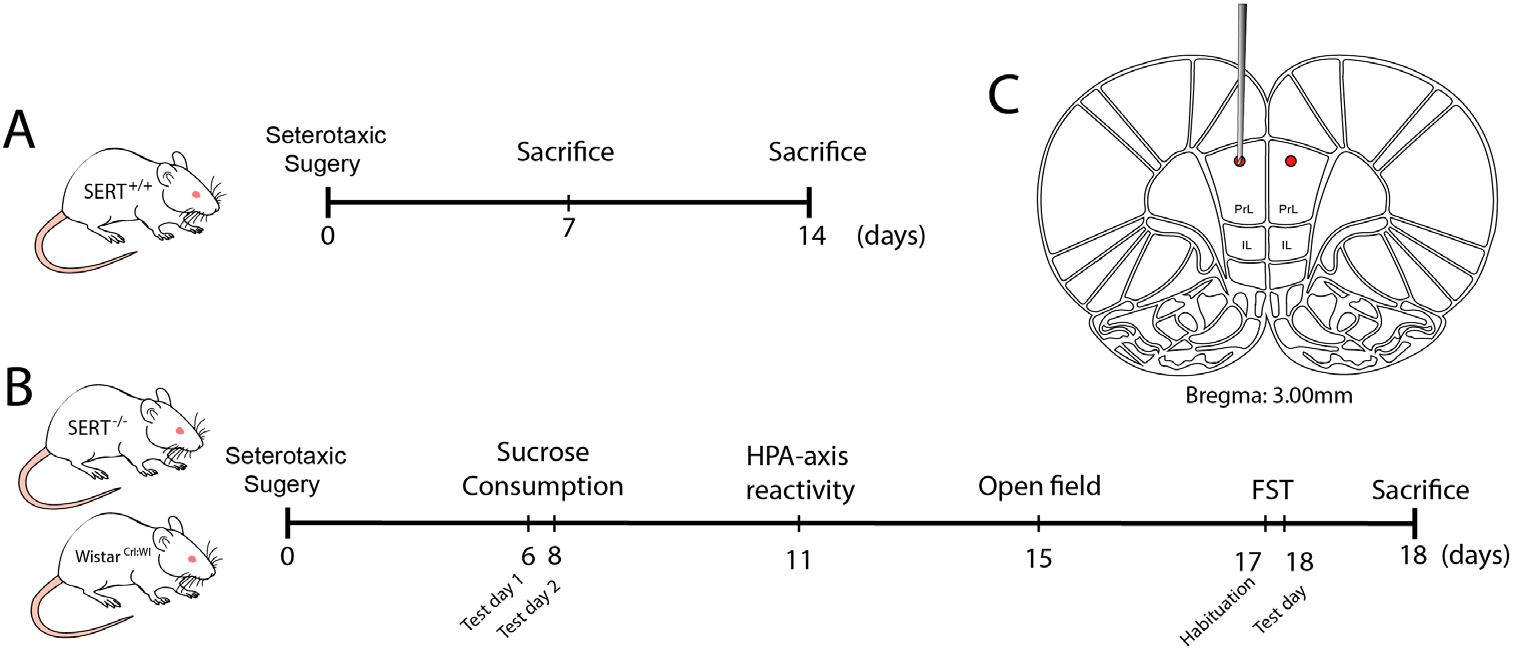
Schematic representation of the experimental design. A) Evaluation of BDNF overexpression in naïve SERT ^+/+^ rats one and two weeks following viral infusion. B) Behavioral tests: Viral infusion followed by behavioral tests including sucrose consumption test, HPA-axis reactivity, open field, and forced swim test (FST). C) Representation of the local o the infusion of either BDNF or control virus.

**Figure 1.**
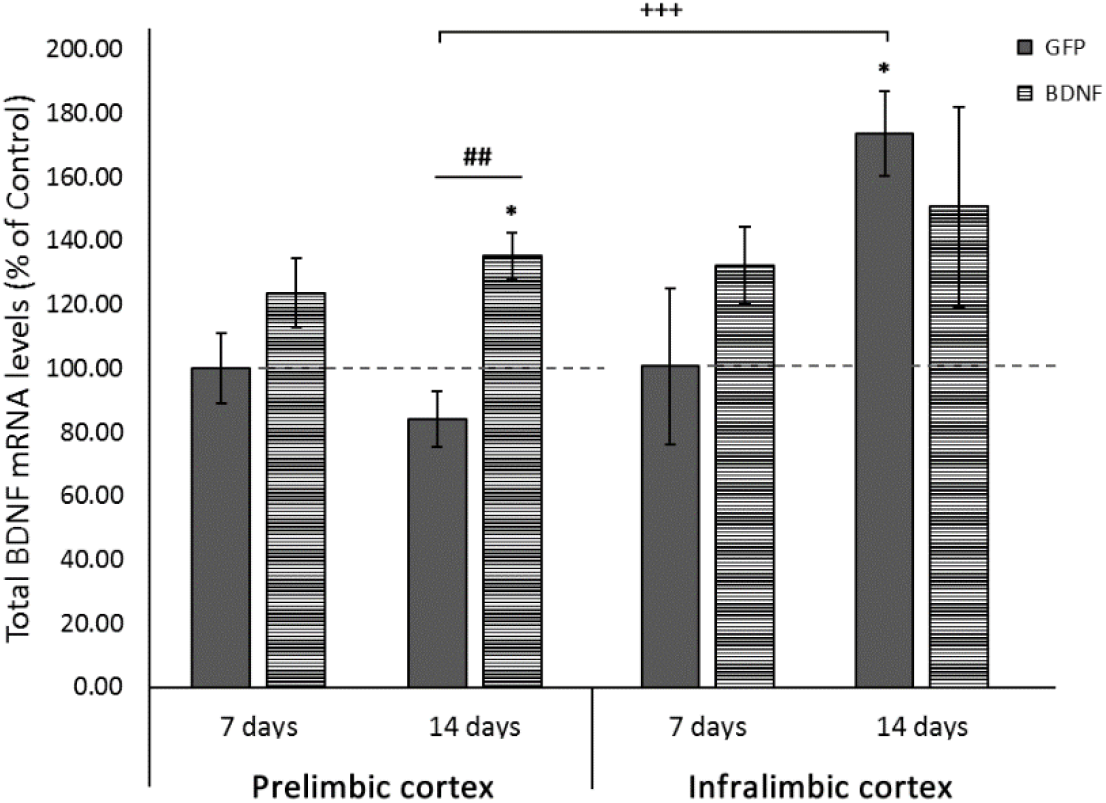
Modulation of total BDNF expression in SERT ^+/+^ animals infused with either GFP control or BDNF viral particles at 1 week and 2 weeks after stereotaxic surgery. Total BDNF mRNA levels were measured in the prelimbic cortex and infralimbic cortex. Data are expressed as fold change compared to the GFP-treated animals (set at 100%), and reflect mean ±SEM from 4-6 independent determinations. * = *p* < 0.05 *vs* GFP-7 days; ## = *p* < 0.01 *vs* GFP-14 days; +++ = *p* < 0.001 IL *vs* PrL (GFP-14 days).

### Stereotaxic Surgery

Rats were anesthetized using isoflurane (5% induction, 2-3% maintenance). Lidocaine (10% m/v) was used for local anesthesia. Animals were fixed in a robot stereotaxic frame (StereoDrive, Neurostar, Germany). The coordinates for the site of the injection were theoretically determined based on the Paxinos & Watson (2007) rat brain atlas and checked through histological evaluation of 30 μm brain slices from dye-infused SERT^+/+^ rats. The total volume of 1μL of either BDNF lentivirus particles (transcript variant IV under CMV promoter, NM_001270633.1) or pLenti-C-mGFP control lentivirus particles, was bilaterally infused into the prelimbic cortex according to the following coordinates: AP +3.0 mm, ML ±0.6 mm, DV −3.0 mm. After surgery, animals were placed in IVC cages (Sealsafe Plus GR900 green line, Tecniplast, Italy) until sacrifice.

### RNA Preparation And Gene Expression Analysis By Quantitative Real-Time PC

Total RNA was isolated from the prelimbic and infralimbic region of the mPFC by single-step guanidinium isothiocyanate/phenol extraction using PureZol RNA isolation reagent (Bio-Rad Laboratories; Segrate, Italy), according to the manufacturer’s instructions, and then quantified by spectrophotometric analysis (NanoDrop™1000, Thermo Scientific). Following total RNA extraction, an aliquot of each sample was treated with DNase to avoid DNA contamination. Then, the samples were processed for real-time PCR to assess total BDNF, BDNF isoform IV, and VI. The analyses were performed by TaqMan qRT–PCR instrument (CFX384 real-time system, Bio-Rad Laboratories S.r.l.) using the iScript one-step RT–PCR kit for probes (Bio-Rad Laboratories). Samples were run in 384-well formats in triplicates as multiplexed reactions with a normalizing internal control (36B4). Thermal cycling was initiated with incubation at 50°C for 10 min (RNA retrotranscription), and then at 95°C for 5 min (TaqMan polymerase activation). After this initial step, 39 cycles of PCR were performed. Each PCR cycle consisted of heating the samples at 95°C for 10 s to enable the melting process and then for 30 s at 60°C for the annealing and extension reactions. Data were analyzed with the comparative threshold cycle (ΔΔCt) method using 36B4 as a reference gene. Primers and probe for BDNF exon IV and VI were purchased from Life technologies (BDNF exon IV: ID EF125679 and BDNF exon VI: ID EF125680). Primers and probe for total BDNF and 36B4 were purchased from Eurofins MWG-Operon. Their sequences are shown below:

- total BDNF: forward primer 5’-AAGTCTGCATTACATTCCTCGA-3’, reverse primer 5’-GTTTTCTGAAAGAGGGACAGTTTAT-3’, probe 5’-TGTGGTTTGTTGCCGTTGCCAAG-3’;
- 36B4: forward primer 5’-TTCCCACTGGCTGAAAAGGT-3’, reverse primer 5’-CGCAGCCGCAAATGC-3’, probe 5’-AAGGCCTTCCTGGCC GATCCATC-3’.

### Behavioral tests

#### Sucrose Consumption Test

After stereotaxic surgery, animals were housed individually and provided with two bottles of water for a 5 days habituation period in which side preference was checked. The sucrose consumption test was adapted from Olivier et al. (2008) and consisted of two days of free-choice access to 24 hours sucrose versus water bottles with a water-only bottle choice in between the two days. In detail, on test-day 1, one of the water bottles was replaced by sucrose 8% solution, and animals had free drinking access for 24 hours. Next, animals received water in both bottles for 24 hours, ending with another 24 hours of free choice between water and sucrose 8% solution on test day 2. The position of the bottles was switched from sucrose consumption test day 1 to the test day 2 to prevent spatial bias. Daily, liquid intake and bodyweight were measured. The data are presented as the preference of sucrose above water (sucrose intake in ml divided by total intake X 100%) and the intake in grams of a 100% sucrose solution per kg bodyweight (intake in ml corrected for the voluminal weight of sucrose and recalculated toward a 100% solution divided by bodyweight in kg).

#### HPA-axis reactivity Test

HPA-axis reactivity was assessed through the measurement of corticosterone levels in the plasma. Usually, when rodents are submitted to stress, plasma concentrations of corticosterone (CORT) peak after 15 to 30 minutes and gradually decrease 60 to 90 minutes later to the pre-stress levels (de Kloet et al., 2005). Therefore, blood samples from tail cuts were collected in capillary blood collection tubes (Microvette^®^ CB 300 Di-Kalium-EDTA, Sarstedt, Germany) 5 minutes before, and 15 and 60 minutes after 30 minutes of restraint stress. Rodent restrainers Broome-style were used for the restraint stress (554-BSRR, Bio-services, The Netherlands). Blood samples were centrifuged (3400 rpm for 15 min at 4 °C), and the plasma was stored at −80 °C until analysis. CORT levels were measured using a radioimmunoassay (RIA) kit according to the manufacturer protocol (ImmuChemTM Double Antibody Corticosterone 125I RIA, MP Biomedicals, USA).

#### Open field test

Novelty-induced locomotor activity was recorded by video recording in Phenotyper^®^ cages (Noldus Information Technology, Wageningen, The Netherlands). The cages (45 cm × 45 cm × 45 cm) were made of transparent Perspex walls and a black floor. Each cage had a top unit containing a built-in digital infrared-sensitive video camera, infrared lighting sources, and hardware needed for video recording. To explore the novelty factor, animals were not exposed to this cage previously and the cages were cleaned with 70% alcohol solution between trials to prevent transmission of olfactory cues. Spontaneous locomotor activity was monitored for 1 hour, and the following parameters were scored using Ethovision XT 11.5 (Noldus Information Technology, Wageningen, Netherlands): distance moved, velocity, frequency and time spent in the center of the cage (Manfré et al., 2017; Schipper et al., 2011a).

#### Forced Swim test

The forced swimming test was performed as previously described (Porsolt et al., 1978). Briefly, rats were individually placed in cylindrical glass tanks (50 cm height, 20 cm diameter) filled to a height of 30 cm with 23±1°C water. The test consisted of two sessions. In the first session, animals were submitted to a habituation period of 15 minutes, then 24 hours later, to a second session of 5 min. The video recordings of the second session were used to automatically score the movements of the rats through a computerized system (Ethovision XT 10, Noldus, The Netherlands). Scored behaviors were ‘immobility’, which reflects no movement at all and/or minor movements necessary to keep the nose above the water; ‘mobility’, indicating movement that corresponds to swimming activity; and ‘strong mobility’, reflecting ‘escape behavior’ (e.g., climbing against the walls and diving). Settings within Ethovision were adjusted based on manually recorded sessions (immobility/mobility threshold: 12; mobility/strong mobility threshold: 16.5 (Boulle et al., 2016; Van den Hove et al., 2013).

### Statistical Analysis

The data were checked for outliers and normality (using the Shapiro–Wilk statistic), and extreme outliers were windsorized. Two-way analysis of variance (ANOVA) was computed for gene expression analysis, with time, genotype, and treatment as independent factors. The outcomes of SPT, Novelty-induced locomotor activity, FST, and post-behavioral gene expression were also analyzed through Two-way ANOVA considering genotype and treatment as fixed factors. Post-hoc Fisher Protected Least Significant Difference (PLSD) or independent sample t-tests were performed where applicable to compare individual group differences. All these statistical analyses were carried out using IBM^®^ SPSS^®^ statistics, version 23 (IBM software, USA). Regarding the HPA-axis reactivity test, a linear mixed model was implemented to account for repeated measurements, and multiple factor analysis using the LME4 package in R (3.5.1). Time, genotype, and treatment effects were modeled as a fixed effect, together with their pairwise double interactions, and their triple interactions. Subject intercepts were modeled as random effects. A likelihood-test ratio was used to assess fixed effect significance. *Post-hoc* tests were performed with the multicomp package, which accounts for multiple hypothesis testing. Significance was accepted at a *p*<0.05 threshold. Descriptive statistics are provided as mean +/− 1 standard error of the mean (SEM).

## Results

### Upregulation of Total BDNF mRNA in naïve SERT^+/+^ rats following prelimbic BDNF lentivirus infusion

Feasibility of BDNF expression and its temporal dynamics was separately examined in a group of naïve SERT^+/+^ rats. mRNA levels were evaluated one and two weeks following BDNF or GFP lentivirus infusion in the prelimbic (PrL) cortex of naïve SERT^+/+^ rats. RT-qPCR was performed to measure total BDNF mRNA overexpression in the prelimbic (PrL) and in in the neighboring mPFC area infralimbic (IL). Two-way ANOVA revealed a significant main effect for treatment in the PrL (F_(1, 18)_ = 13.790, *p* = 0.002). PLSD *post-hoc* analysis revealed BDNF overexpression in the site of the injection (PrL) with significant total BDNF mRNA increased in the BDNF treated animals compared to the control GFP treated rats both one week (*p* = 0.038) and two weeks (*p* = 0.002) after surgery. Interestingly, while in the PrL, no time point differences were identified among control GFP treated rats, the IL samples presented a sharp rise (72.47 %, SD = 26.41) in BDNF levels in control-treated animals (*p* = 0.01). As shown in figure 2, this increase led to significantly higher BDNF levels in the IL than in the PL (*p* < 0.001). We concluded that BDNF overexpression was stable for at least 14 days, specifically in the PrL.

**Figure 2.**
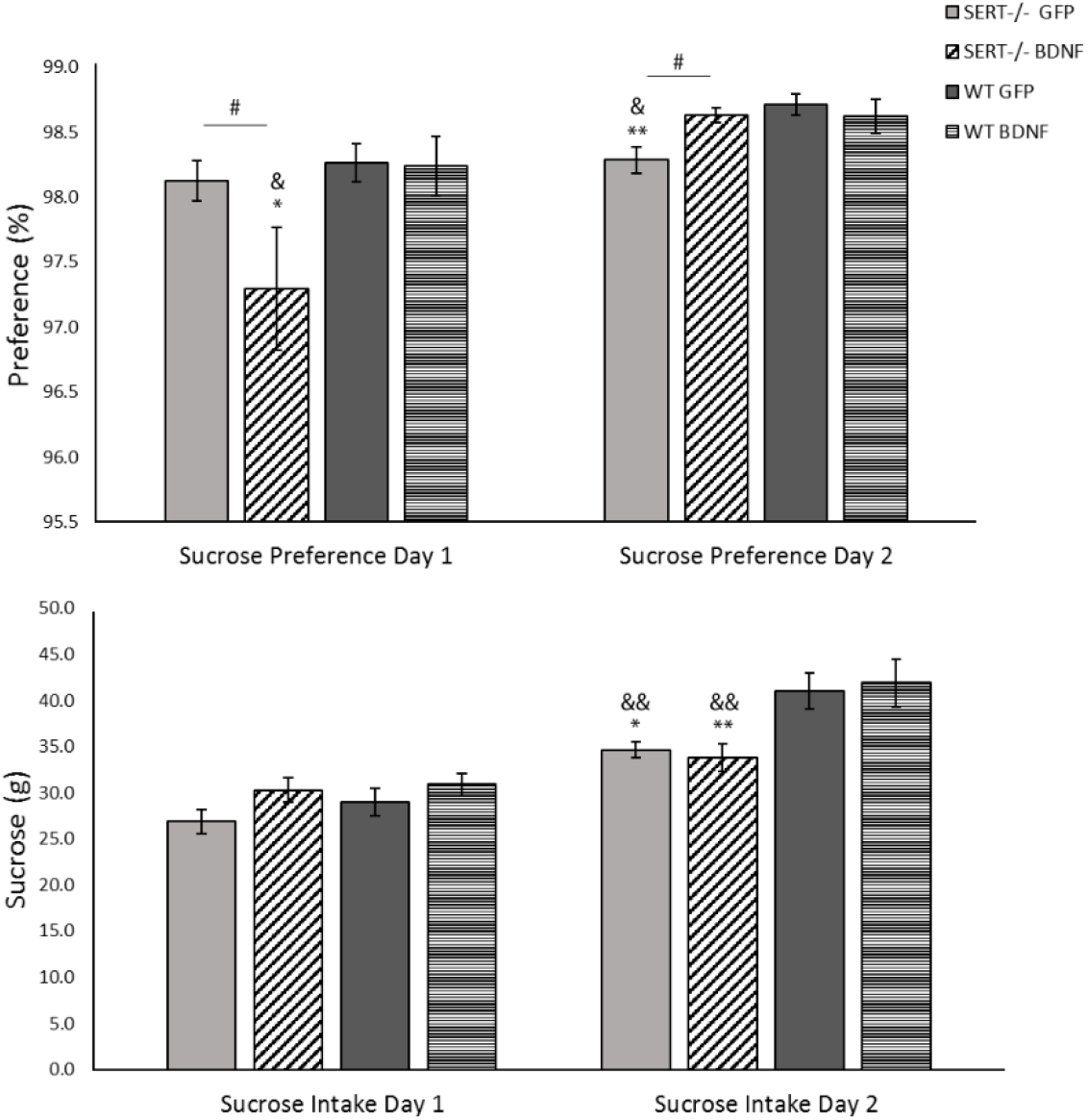
Sucrose consumption of 8% sucrose solution by SERT^−/−^ and WT rats. Data are expressed as mean S.E.M. sucrose preference (sucrose intake/total fluid intake x 100%), and as mean S.E.M. total sucrose intake (g) per body weight (n = 10-12; * = p < 0.05 and ** = p < 0.01 vs WT-GFP; & = p < 0.05 and && = p < 0.01 vs WT-BDNF; # = p < 0.05 vs SERT^−/−^GFP).

### Sucrose Consumption Test (SCT)

Anhedonia is marked by a reduced interest in pleasurable events, and it is present in depression. This depression-like symptom can be identified in rodents through a decrease in sucrose consumption. Animals were exposed to two days of free access to sucrose 8% solution. The results of the sucrose intake in grams and the preference for sucrose above the water are described below.

#### Sucrose Preference in SERT^−/−^ rats is altered by BDNF overexpression

On the first day of testing, no significant main effects were observed for sucrose preference. Pairwise comparisons, however, revealed that sucrose preference was significantly reduced in the SERT^−/−^ rats treated with BDNF lentivirus compared to untreated SERT^−/−^ (*p* = 0.05) and control WT rats (*p* = 0.018). At the second day of testing, two-way ANOVA analysis showed a genotype as well as a genotype versus treatment interaction (F_(1, 40)_ = 4.738, *p* = 0.035 and F_(1, 40)_ = 5.058, *p* = 0.030, respectively). As seen in figure 3, the *post-hoc* analysis further demonstrated that SERT^−/−^ rats treated with BDNF presented a higher preference for sucrose than control-treated SERT^−/−^ animals (*p* = 0.017). Additionally, control-treated SERT^−/−^ animals displayed lower sucrose preference than the control WT rats (*p* = 0.002). Therefore, in summary, BDNF treatment in SERT^−/−^ rats improved the preference for sucrose in the second day of the test.

**Figure 3.**
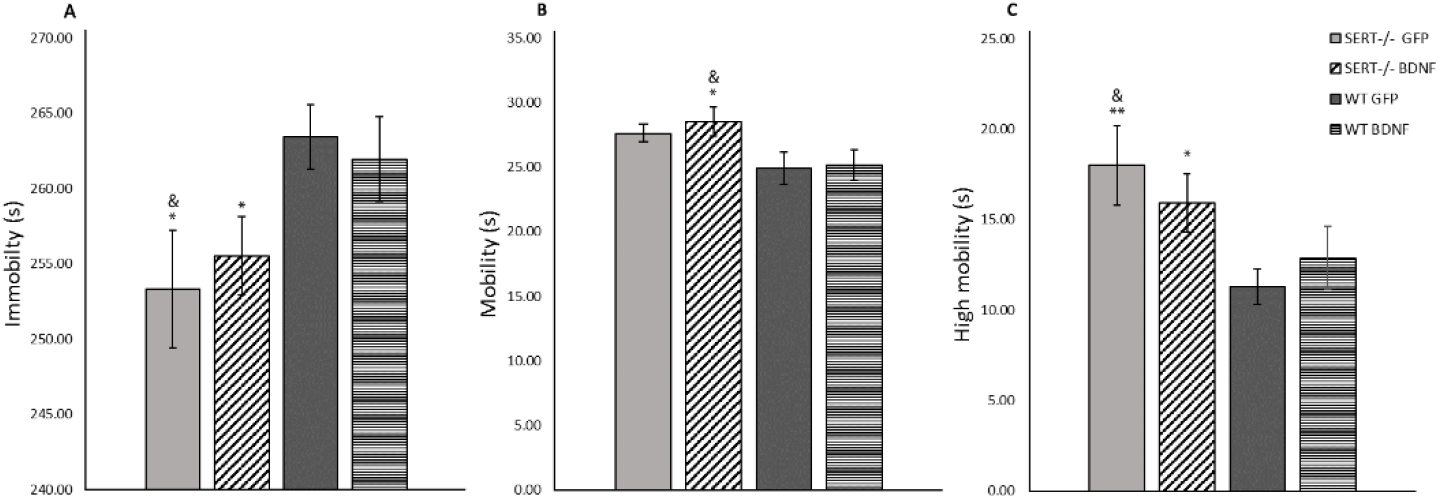
Mean (±SEM) measure of (A) immobility, (B) mobility, and (C) high mobility in the forced swim test. n = 10–12 rats per group. * p < 0.05 and ** p < 0.01 vs WT GFP, & p <0.001 vs WT BDNF). Two-way ANOVA, Fisher LSD post-hoc test.

#### SERT^−/−^ genotype rather than BDNF overexpression modulates the rat’s response to sucrose intake

As for sucrose preference, two-way ANOVA did not reveal a significant main effect for sucrose intake in SERT^−/−^ versus WT rats on the first day of the test. There were also no statistically significant differences in the amount of sucrose consumed among the groups (Figure 3). On the second day of testing, however, a main genotype effect was found for sucrose consumption (F_(1, 42)_ = 16.789, *p* < 0.001). Moreover, *post-hoc* analysis showed that SERT^−/−^ rats consumed significantly less sucrose than controls (*p* < 0.05) with no treatment differences. We concluded that as previously described (Olivier et al., 2008), the SERT^−/−^ phenotype led to reduced sucrose consumption that was not rescued by the PrL BDNF transfection.

### BDNF overexpression did not modify SERT^−/−^ rats anxiety-like pattern in the Forced Swim Test

When rodents are exposed to an inescapable stressor such as in the forced swim test, their motivation to cope with stress can be quantified by the percentage of time spent on immobility (behavioral passivity) or performing a highly mobile (escape-like) behavior (Porsolt et al. 1977). We found a main genotype effect for immobility (F_(1,42)_ = 7.827, *p* = 0.008). Unexpectedly, as figure 4 shows, immobility was decreased in both control- and BDNF-treated SERT ^−/−^ rats compared to WT rats (*p* = 0.017 and *p* = 0.027, respectively). Additionally, two-way ANOVA revealed a genotype main effect for high mobility (F_(1, 42)_ = 8.278, *p* = 0.006). Particularly, *post-hoc* examination demonstrated that the time spent on high mobility swimming or escape behavior was significantly higher in SERT^−/−^ rats than in WT controls (*vs.* SERT^−/−^ GFP *p* = 0.006, *vs.* SERT^−/−^ BDNF *p* = 0.021). In summary, surprisingly, SERT^−/−^ rats presented decreased immobility and increased escape behavior compared to WT animals.

**Figure 4.**
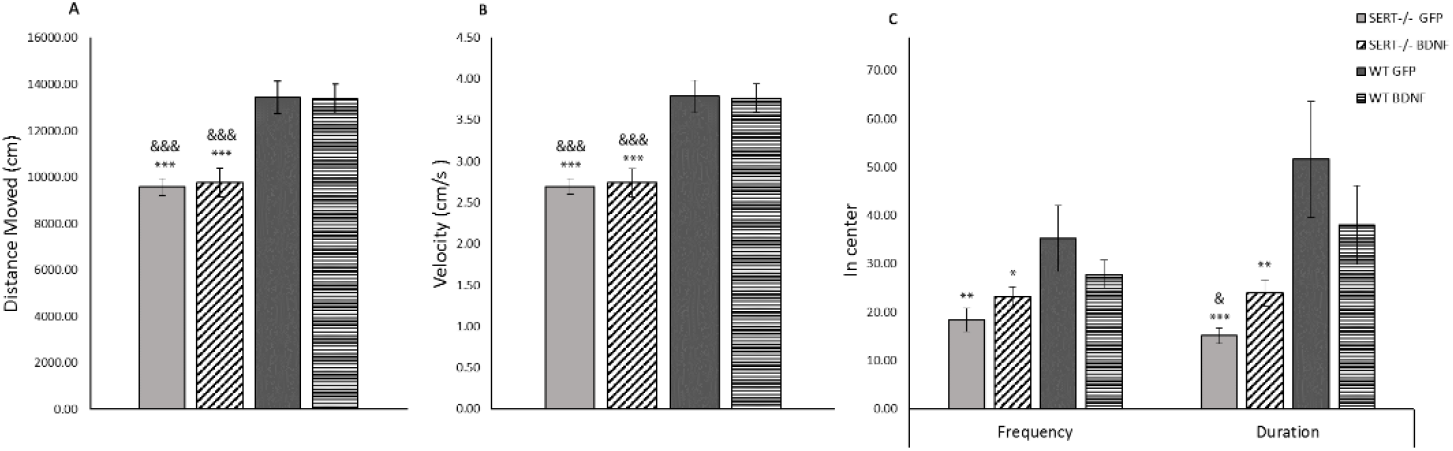
Novelty-induced locomotor activity expressed as mean (±SEM) measure of (A) distance moved, (B) velocity and (C) frequency and time spent in center. n = 10-12; * = p < 0.05, ** = p < 0.01, *** = p < 0.001 vs WT GFP; & = p < 0.05 and &&& = p < 0.001 vs WT BDNF.

### Novelty-induced locomotor activity is impaired in SERT^−/−^ rats

Rodents may present higher activity when they are introduced to a novel environment (Menzaghi et al., 1994). A decrease in central locomotion (frequency and time spent in the central part of the arena), together with a general decrease in the locomotion (distance moved and velocity) can be interpreted as an anxiogenic-like behavior (Prut and Belzung, 2003). In our experimental conditions, genotype played a role in the locomotor activity affecting distance moved (F_(1,42)_ = 40.708, *p* < 0.001), velocity (F_(1,42)_ = 41.559, *p* < 0.001), time spent in the center of the arena, as well as the frequency to which the animals accessed the center (F_(1, 42)_ = 11.621, *p* = 0.001) and F_(1, 42)_ = 6.743, *p* < 0.013, respectively). No treatment effect was found for SERT^−/−^ or WT rats exposed to the new environment. Further *post-hoc* investigation revealed that SERT^−/−^ rats, independently of treatment, displayed a significant decrease in distance moved and velocity (*p* < 0.0001 for all SERT^−/−^ *vs* WT rats comparisons); additionally, as it can be seen in figure 5, frequency and duration spent in the center of the arena were reduced in SERT^−/−^ BNDF-treated rats (frequency: *vs* WT GFP *p* = 0.04; duration: *vs* WT GFP *p* = 0.01) and in SERT^−/−^ GFP treated rats (frequency: *vs* WT GFP *p* = 0.005; duration: *vs* WT GFP *p* < 0.001, *vs* WT BDNF *p* = 0.039). In brief, exposition to a novel environment caused an anxiety-like response in SERT^−/−^ rats as expressed by decreased mobility and decreased access to the center.

**Figure 5.**
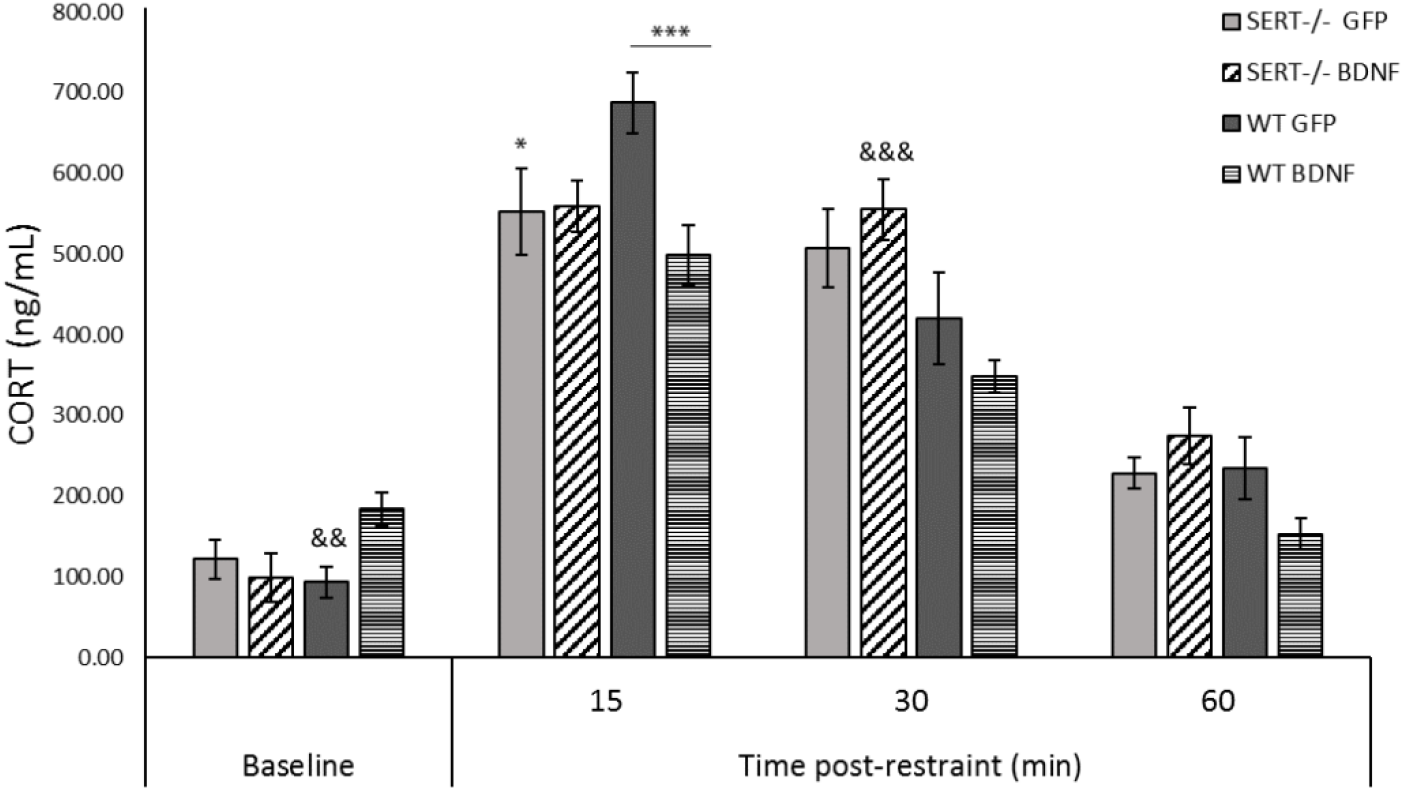
HPA-axis reactivity assessment. Corticosterone (CORT) levels are expressed mean ±1 standard error of the mean (SEM) of measurements from plasma samples obtained 5 minutes before restraint stress (baseline), and 15, 30, and 60 minutes post-restraint stress. n = 10-12; * = p < 0.05, *** = p < 0.001 vs WT GFP; && = p < 0.01 and &&& = p < 0.001 vs WT BDNF. Linear mixed model.

### BDNF overexpression exclusively alters HPA-axis reactivity in WT animals

Acute restraint stress can activate the hypothalamic-pituitary-adrenal axis (HPA-axis), resulting in the release of CORT in rodents, which can be a measure of the (mal)functionality of the HPA-axis. The stress-induced increase in CORT levels relative to baseline was evaluated 15, 30, and 60 minutes after stress. A linear mixed-effect analysis indicated a triple interaction effect between treatment, genotype, and time (F_(15,18)_ = 15.84, p = 0.001). A post-hoc analysis comparing groups pairwise per time point indicated a strong difference between BDNF vs. GFP transfected WT rats at the 15 min time point (p < 0.001) and between WT and SERT^−/−^ rats in the GFP condition (p < 0.03). This difference indicates that BDNF overexpression in the PrL has the potential to reduce the HPA reactivity during the first phase of the response, although only in the WT group. Interestingly, CORT baseline levels were increased in BDNF-treated WT animals (t-test *vs.* WT GFP *p* = 0.003). At the 30 min time point, a difference between BDNF-transfected WT and SERT^−/−^ rats was also found (*p* < 0.001), indicative of an elevated CORT level in SERT rats relative to WT control. All but the WT BDNF group after 60 min still displayed CORT levels significantly elevated from baseline, indicative of a strong activation of the HPA-axis, with a trend toward return near baseline after 60 min. In summary, BDNF overexpression appeared to increase basal CORT levels and decrease the HPA reactivity in the WT group, whereas the SERT^−/−^ rats were unaffected.

### BDNF overexpression was stable in WT animals after behavioral challenge

To assess the BDNF expression induced by transfection in the behaviorally tested animals, rats were sacrificed 90 minutes after the forced swim test. Overexpression of total, exon IV, and exon VI BDNF transcripts in the PrL and IL were evaluated through RT-qPCR. No main effects were found for total BDNF or BDNF exons IV and VI in the PrL. Pairwise comparisons were computed and, as it is demonstrated in figure 7, they revealed that WT rats treated with BDNF presented higher BDNF levels than GFP-treated SERT^−/−^ rats (total BDNF: *p* = 0.04, BDNF IV: *p* = 0.039, and BDNF VI: *p* = 0.05), as well as higher BDNF levels than BDNF-treated SERT^−/−^ rats (BDNF IV: *p* = 0.016, and BDNF VI: *p* = 0.015).

In the IL, on the other hand, BDNF expression was decreased in both SERT^−/−^ groups compared with both WT groups. As a result, a significant genotype main effect was observed for total BDNF (F_(1, 17)_ = 22.365, *p* < 0.001), BDNF IV (F_(1, 18)_ = 9.707, *p* = 0.006), and BDNF VI (F_(1, 17)_ = 13.397, *p* = 0.002). Post-hoc testing demonstrated no treatment differences, but did reveal a significant BDNF downregulation of total, exon IV, and exon VI BNDF transcript levels in SERT^−/−^ rats compared to WT rat controls (*p* < 0.05). In conclusion, we observed that BDNF overexpression in the PrL of WT animals remained even after the behavioral tests; moreover, in SERT^−/−^ rats no changes were found in BDNF expression levels in the PrL area compared to WT rats, but we did find a decrease in BDNF expression in the IL.

**Figure 6.**
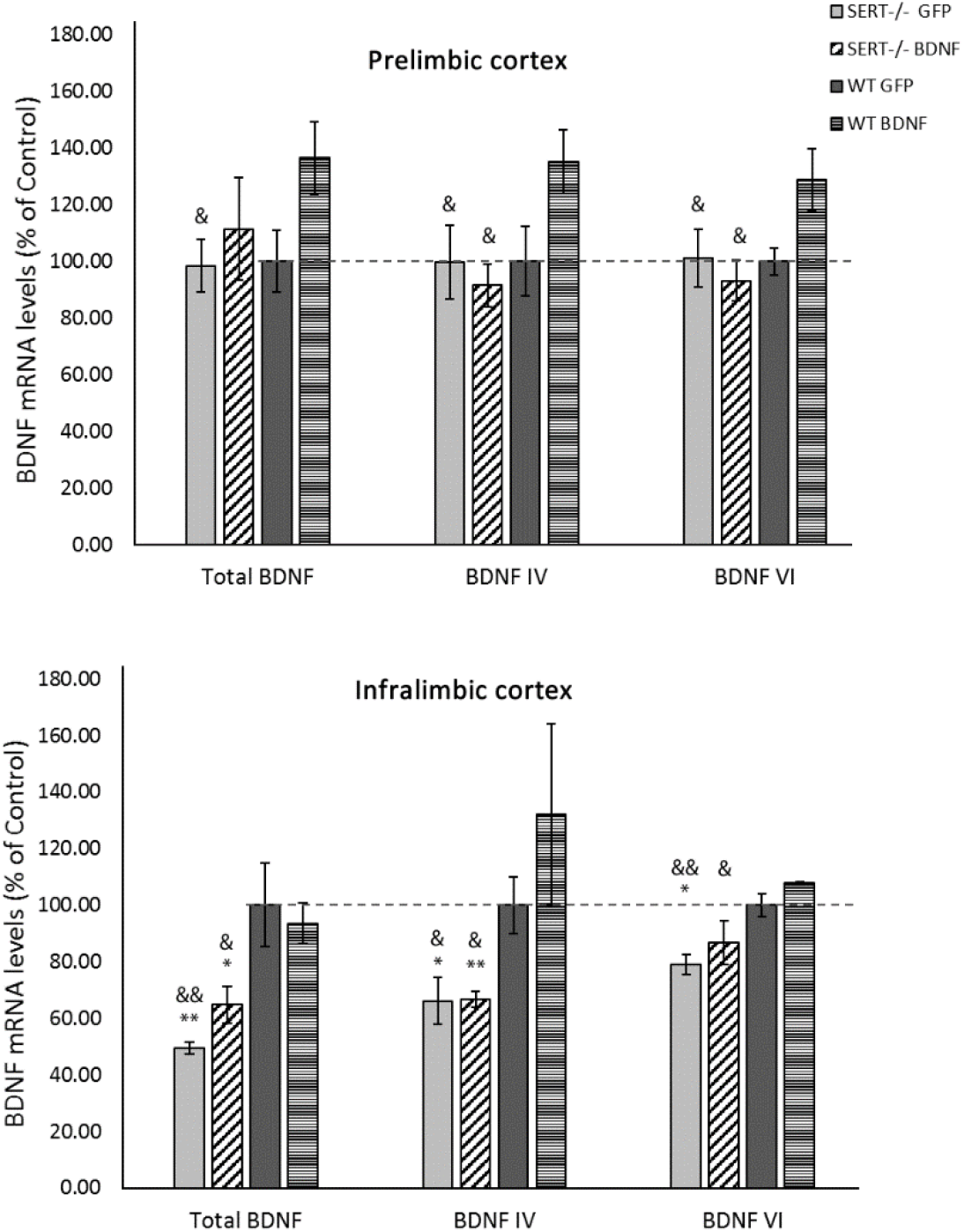
BDNF expression of total, exon IV, and exon VI transcripts in SERT +/+ and SERT KO animals submitted to local infusion of either GFP control or BDNF viral particles followed by sequential behavioral challenges. Total BDNF, BDNF IV, and BDNF VI mRNA levels were measured in the prelimbic cortex and infralimbic cortex. Data are expressed as percentage change compared to the WT GFP-treated animals (set at 100%), and reflect mean ±SEM from 4-6 independent determinations. * = p < 0.05 and ** = p < 0.01 vs WT GFP; & = p < 0.05 and && = p < 0.01 vs WT BDNF.

## Discussion

In this study, we have demonstrated that BDNF IV lentiviral infusion into the PrL induced overexpression of total BDNF in naïve SERT^+/+^ animals in a time- and brain region-dependent manner. Notably, we confirmed increased mRNA BDNF levels in the PrL of BDNF lentivirus treated SERT^+/+^ rats two weeks after the surgery, as well as an unexpected BDNF upregulation in the IL of control-treated animals. Moreover, BDNF overexpression caused different phenotypical outcomes depending on the behavioral task and the animal genotype. In WT animals, BDNF upregulation caused an alteration in the HPA-axis function following acute restraint stress, and in SERT^−/−^ rats, it resulted in improvement of the anhedonia-like symptoms in the sucrose preference test. Whereas the stress coping behavior as measured in the forced swim test and locomotor activity were not affected by BDNF overexpression, rather, the SERT^−/−^ genotype played a role in inducing anxiety-like phenotypes in these tests.

During the sucrose consumption test, SERT^−/−^ and WT rats were submitted to two sessions of 24 hours exposition to an 8% sucrose solution. In consequence, we observed that SERT^−/−^ treated with control virus did not differ in sucrose preference from WT rats on the first day of testing, but presented decreased preference in the second day. This result is in line with our previous demonstration that preference for sucrose is negatively affected in SERT^−/−^ animals (Olivier et al., 2008). The differences between the first and second day of sucrose preference seen in SERT^−/−^ rats have been demonstrated before in studies using a mouse animal model presenting selective disruption of BDNF IV (BDNF-KIV mice) in which BDNF-KIV mice only exhibited significantly decreased sucrose preference in the second day of sucrose test (Sakata et al., 2010). Accordingly, SERT^−/−^ rats treated with BDNF IV lentivirus presented reduced sucrose preference in the first day, but increased preference in the second day when compared to control-treated SERT^−/−^ and WT rats. Therefore, it is likely that BDNF IV overexpression in SERT^−/−^ rats led to neophobia to a novel taste or an anxiety-like behavior upon the first exposure; while during the second exposure, when the animals were familiar with the new taste, the anhedonia-like behavior in SERT^−/−^ animals was suppressed by the BDNF overexpression. Conclusively, possibly, overexpression of BDNF transcript IV in the PrL was responsible for rescuing the anhedonia-like behavior displayed by SERT^−/−^ rats.

Concerning the forced swim test, no BDNF treatment differences were found, but we did notice apparent genotype behavioral differences. In contrast to previous studies employing SERT^−/−^ rats and mice, SERT^−/−^ rats spent less, rather than more, time on immobility than WT animals (Lira et al., 2003; Olivier et al., 2008). In agreement with our current results, studies in other animal models presenting alterations in the BDNF system have also shown that subjects with decreased BDNF levels did not display higher immobility in the forced swim test compared to controls (Duman and Monteggia, 2006; Sakata et al., 2010). In our experimental conditions, decreased immobility can be justified by the fact that SERT^−/−^ rats spent proportionally more time on vigorously strong swimming (high mobility) than immobility compared to the WT animals. Several lines of research have hypothesized that this escape behavior essentially reveals an increased level of anxiety (Anyan and Amir, 2018). As mentioned before, SERT^−/−^ animals have impaired BDNF expression as well as reduced expression of its transcription factors such as Npas4. Both BDNF and Npas 4, are implicated in the establishment of the GABAergic system. The GABAergic system communicates with other neurotransmitter networks, and its downregulation can lead to an anxiety-like behavioral outcome (Lydiard, 2003; Millan, 2003). Correspondingly, SERT^−/−^ rats present changes in the functioning of the GABAergic system (Calabrese et al., 2013; Luoni et al., 2013; Miceli et al., 2017; Schipper et al., 2019). Therefore, likely, reduced levels of GABA in these animals could contribute to an anxiety-like response upon behavioral challenges such as the forced swim test.

In the novelty-induced locomotor activity test, the SERT^−/−^ rats presented decreased locomotor activity in a novel environment compared to the WT animals. This decrease indicates that the reduced immobility and increased high mobility observed in SERT^−/−^ rats in the forced swim test was not due to increased locomotor activity. Moreover, similarly to the forced swim test, the decreased locomotor activity seen in the SERT^−/−^ rats contradicts previous studies using SERT^−/−^ rats, which report no genotype differences between naïve SERT^−/−^ and SERT^+/+^ rats in the novelty-induced locomotor activity test (Homberg et al. 2008; Schipper et al. 2011). However, in contrast to the findings in rats, studies in SERT^−/−^ mice have shown that in an unfamiliar environment, SERT^−/−^ mice displayed reduced locomotor activity and anxiety-like behavior similar to our results (Alexandre, 2006; Kalueff et al., 2007). Hence, the reduced distance moved, lower velocity, and especially the decreased time and frequency in the center of the arena might be a reflection of the overall increased anxiety-like behavior displayed by SERT^−/−^ rats which have also been described in SERT^−/−^ mice (Olivier et al., 2008).

We further evaluated the HPA-axis activity to assess the effects of BDNF overexpression in SERT^−/−^ and WT rats. Our results revealed that basal levels of CORT were not altered in SERT^−/−^ rats compared to control-treated WT rats. Conversely, we found that BDNF upregulation in WT rats generated increased basal levels of CORT in this group when compared to control WT animals. Moreover, after acute restraint stress, WT rats treated with BDNF presented similar HPA-axis response to the SERT^−/−^ animals, namely a decreased elevation in CORT levels compared to control-treated WT rats. Therefore, these results show that BDNF upregulation altered the HPA-axis function in WT rats with no effects in SERT^−/−^ animals. Concerning the HPA-axis disturbances seen in WT rats overexpressing BDNF, likely basal HPA-axis hyperactivity and decreased response to stress might have been facilitated by a discrepancy between the rate of mature BDNF (mBDNF) protein to its precursor proBDNF, favoring the later one. This imbalance in the proBDNF/mBDNF was confirmed before in a study showing that BDNF overexpression can lead to an increase in the release of uncleaved proBDNF (Leßmann and Brigadski, 2009). While mBDNF supports plasticity through its high affinity for the TrkB receptor, proBDNF has an affinity for the p75NTR receptor, which mediates apoptotic signaling leading to neuronal death (Woo et al. 2005; Leßmann and Brigadski 2009). Although both HPA hyperactivity and increased proBDNF are present in depressive disorders (Bai et al. 2016; Zhou et al. 2013; Arborelius et al. 1999; Nestler et al. 2002; Pariante and Lightman 2008; van Bodegom, Homberg, and Henckens 2017), the direct proof that the deleterious effects of proBDNF are the underlying cause to HPA-axis malfunction is still lacking. Yet another possibility we tend to support to explain the disruption in the HPA-axis reactivity in WT rats overexpressing BDNF is that the gene upregulation led to exceeding levels of the mature form of the BDNF protein. Accordingly, the exceeding mBDNF in the PrL would preferentially inhibit CORT production at the amygdala level, shortening the HPA-axis response loop, resulting in decreased CORT release upon stress in comparison to control animals. The PFC, which is one of the key regions in the control of the HPA-axis, presents inhibitory connections with the amygdala and PVN (van Bodegom et al., 2017), and it has been demonstrated that when BDNF is overexpressed in the PFC, it can undergo anterograde transport and cause BDNF overexpression in the amygdala (McGinty et al., 2010). BDNF, especially exon IV, is known for its critical role in GABAergic transmission (Sakata et al., 2009). Brivio et al. (2020) showed that acute restraint stress increased PFC levels of total and BDNF exon IV in Sprague Dawley rats specially 1 hour following the acute stress. Total BDNF levels were also increased in the PFC following acute swim stress (Brivio et al., 2019). Therefore, acute restraint stress may have caused activity-dependent upregulation of BDNF (Leßmann and Brigadski, 2009), leading to enhancement in the GABAergic inhibitory control in the amygdala, and consequently, increased negative feedback to the HPA-axis system (Barry et al., 2017; Liu et al., 2014; Zhang et al., 2018).

In conformity, because BDNF IV, the transcript chosen to be upregulated in the present study, is activity-dependently released (Leßmann and Brigadski, 2009), we also evaluated the BDNF mRNA levels in the PrL and IL of the animals submitted to behavioral testing. Considering the intricate control over the BDNF gene, in which different isoforms are generated by distinct promoters (Aid et al., 2007), and giving that different stimuli can influence the isoforms response and cellular location, we focused not only on total mRNA but also BDNF exon IV and VI transcripts. These transcripts are involved in depressive- or anxiety-like behavior and are known to be downregulated in SERT^−/−^ rats (Molteni et al., 2010; Sakata et al., 2010). As a result, in the PrL, we did observe that WT rats receiving BDNF lentivirus presented a higher BDNF expression than the WT control group and SERT^−/−^ rats. However, when compared to the control WT group, this overexpression was not statistically significant. Furthermore, we did not observe any differences in the level of all analyzed BDNF transcripts when comparing both SERT^−/−^ groups to control WT rats. This finding indicates that BDNF overexpression did not change levels of BDNF in SERT^−/−^ rats compared to both control-treated SERT^−/−^ and WT rats. However, taking into consideration that the gene expression analysis was conducted after the behavioral testing, possibly the viral transfection in the PrL was not stable after exposing the animals to behavior challenges. Additionally, we noticed that when comparing control-treated SERT^−/−^ and WT no differences in BDNF levels were found in the PrL. Conversely, in the IL, a remarkable downregulation in total BDNF, BDNF IV and BDNF VI was seen in the SERT^−/−^ animals in comparison to WT controls. The observation that mRNA BDNF levels were unchanged in SERT^−/−^ even after viral upregulation may help to understand the mechanisms behind the outcomes seen in the behavioral tests. For instance, we have demonstrated that SERT^−/−^ rats displayed anxiety-like behavior, presenting higher activity in the forced swim test, less activity in the novelty-induced locomotor activity, and altered HPA-axis response upon restraint acute stress. A possible explanation for the overall anxiety-like behavior in SERT^−/−^ rats may be based on the complex control the BDNF gene can undergo. For example, previous research showed that transcription factors that regulate the BNDF transcription, such as CREB, Arnt2, CaRF, NFkB, and Npas4, are significantly downregulated in SERT KO rats. Changes in Npas4 are directly correlated with decreased BDNF exons I and IV, bringing about the hypothesis that behavioral outcomes related to SERT knockout and BDNF downregulation in the SERT^−/−^ rats might be, at least in part, also attributed to Npas4 downregulation (Guidotti et al., 2012). Therefore, it is appropriate to infer that overexpression of exogenous BDNF IV transcript in the PrL was affected by the downregulation of endogenous transcription factors in the SERT^−/−^ rats. Thus, despite gene overexpression, BDNF protein levels may not have changed in the SERT^−/−^ rats, explaining the lack of behavioral changes upon BDNF overexpression in these animals and its general anxiety-like behavior. On the other hand, this hypothesis does not justify the results observed in the sucrose preference test, where we revealed melioration in the anhedonia-like symptoms of SERT^−/−^ rats treated with BDNF lentivirus. Therefore, because in the present study we did not investigate the levels of BDNF protein in the PrL, possibly other molecular mechanisms might be involved in the BDNF gene regulation concerning the anhedonia-like behavior.

In conclusion, we have shown that BDNF overexpression in the PrL, in general, did not rescue SERT^−/−^ from its depression- and anxiety-like behavior, as demonstrated by the decreased sucrose intake, reduced locomotor activity, and increased high mobility in the forced swim test compared to controls. However, in the sucrose preference test, SERT^−/−^ rats treated with BDNF IV lentivirus presented a higher preference for sucrose (that is a reduction in anhedonia-like behavior) than control SERT^−/−^ animals. Furthermore, BDNF upregulation in WT rats specifically promoted alteration in the HPA-axis activity of WT rats, resulting in increased basal levels of CORT and making these rats respond similarly to the SERT^−/−^ rats upon restraint stress.

This study, however, presented several limitations. For example, the overexpression of an individual transcript variant, namely BDNF IV, in the PrL may have resulted in activation and reinforcement of particular neural networks. These networks could mainly cause HPA-axis alterations in WT animals and reduction of anhedonia-like behavior in SERT^−/−^ rats, without affecting other relevant neuronal circuits involved in the behavioral challenges the animals were submitted. Moreover, although our SERT^+/+^ and SERT^−/−^ have a Wistar background, the outcrossing may not eliminate all additional induced mutations. Therefore, the use of commercial wild type Wistar rats as controls may pose a disadvantage when relating to our previous findings. Nevertheless, the stability of the SERT^−/−^ phenotype across studies, generations, and laboratories have shown that this animal model is a useful tool for studying the effects of life-time increased extracellular serotonin and downregulated BDNF levels (Homberg et al., 2014; Olivier et al., 2010).

Despite the progress in the understanding of the biological effects of BDNF, important aspects of the BDNF gene regulation, as well as the spatiotemporal release and the precise sites of the BDNF action are still poorly understood, especially in the prefrontal cortex (Sakata et al., 2009). Different stimuli seem to differently regulate the transcription of BDNF in specific brain areas adding complexity to the study of the mechanisms behind the effects of BDNF (Adachi, 2014; Baj et al., 2011; Govindarajan et al., 2006; Maynard et al., 2016). The neurotrophic hypothesis of depression was developed based on the observation that stress, anxiety, and depression are accompanied by decreased levels of BDNF, and that several treatments used for such disorders also increase BDNF levels (Duman and Monteggia, 2006). Since vulnerability to depression can be attributed to poor neuronal plasticity (McClung and Nestler, 2008) and underlying neurobiological processes might be associated with BDNF levels, it is likely that changes in BDNF may contribute to an improvement of behavioral symptoms in depressive individuals. Furthermore, taking into account that some of the current first-line treatments for depression targeting the serotoninergic system have failed to work consistently in all patients (Cuijpers, 2017), BDNF is placed as an important candidate for therapeutic modulation in mood disorders in humans. From this perspective, since our study has shown that therapeutic approaches aiming BDNF overexpression may need to be specific to promote symptoms attenuation, it is essential to elucidate further the relevance of the BDNF downregulation found in the PFC or other brain areas of SERT^−/−^ rats as regarding to the neuropathology of depression.

